# Extracellular vesicle fusion visualized by cryo-EM

**DOI:** 10.1101/2022.03.28.486013

**Authors:** Mattia I. Morandi, Petro Busko, Efrat Ozer-Partuk, Suman Khan, Giulia Zarfati, Yael Elbaz-Alon, Paula Abou Karam, Tina Napso Shogan, Lana Ginini, Ziv Gil, Neta Regev-Rudzki, Ori Avinoam

## Abstract

Extracellular vesicles (EVs) transfer bioactive molecules between cells in a process reminiscent of enveloped viruses. EV cargo delivery is thought to occur by protein-mediated and pH-dependent membrane fusion of the EV and the cellular membrane. However, there is a lack of methods to identify the fusion proteins and resolve their mechanism. We developed and benchmarked an *in vitro* biophysical assay to investigate EV membrane fusion. The assay was standardized by directly comparing EV- and viral-fusion with liposomes. We show that EVs and retroviruses fuse with liposomes mimicking the membrane composition of the late endosome in a pH and protein-dependent manner. Moreover, we directly visualize the stages of membrane fusion using cryo-electron tomography. We find that, unlike most retroviruses, EVs remain fusogenic after acidification and re-neutralization. These results provide novel insights into the EV cargo delivery mechanism and an experimental approach to identify the EV fusion machinery.

## Introduction

Extracellular vesicles (EVs) are membrane-enclosed compartments ranging from 50 to 500 nm in diameter loaded with proteins, lipids, RNA, and DNA. They are secreted from several cell types and generally promote physiological and pathological processes, including the immune response, cancer development and metastasis (*1–6*). They are also extensively studied for their potential clinical application as diagnostic biomarkers and drug delivery systems (*3, 7, 8*).

EVs have been classified into three major subpopulations (i.e., microvesicles, apoptotic bodies, and exosomes), each composed of a heterogeneous pool of vesicles (*9*). While microvesicles and apoptotic bodies bud from the plasma membrane, exosomes bud into the lumen of multivesicular bodies (MVBs) and exit cells after MVB fusion with the cell membrane (i.e. MVB exocytosis) (*10, 11*). Regardless of their classification, EVs enter recipient cells from the extracellular environment primarily through vesicular uptake and must release their cargo into the cytoplasm to modulate cell physiology (*12*).

The biogenesis, size, and composition of EVs are remarkably similar to many enveloped single-stranded RNA viruses such as Rhabdoviruses (e.g., Vesicular Stomatitis Virus), Orthomyxoviruses (e.g., Influenza), and Retroviruses (e.g., Human Immunodeficiency Virus) (*13, 14*). The envelopes of such viruses contain glycoproteins that function as ligands to attach the virus to specific cellular receptors and then mediate fusion between the cell membrane and the viral envelope (*15*). These glycoproteins are also frequently essential for virion assembly and budding (*16*). As such, membrane fusion has been suggested as the primary mechanism of EV cargo delivery (*17–20*).

EV cargo delivery has been shown to depend on proteins (*19, 20*) and to be triggered by low pH (*17, 19, 20*). Moreover, the efficiency of cargo delivery can be modulated by changing the lipid composition in the endosome (*17, 18*). These findings reinforce the hypothesis that most EV cargo delivery occurs by EV membrane fusion, triggered by the late endosomal milieu. Nevertheless, the fusion mechanism remains incompletely understood and it is unclear how triggering fusion by low pH is compatible with exosome biogenesis in the acidic MVBs (*21*).

Based on the biophysical assays developed to study viral fusion, we developed an *in vitro* assay to investigate EV membrane fusion by probing the interaction of EVs with artificial liposomes. Using a Forster Resonance Energy Transfer (FRET)-based assay to directly compare EV and viral fusion at the level of membrane mixing *in vitro*, we demonstrate that EVs can fuse unilaterally to lipid membranes in a pH-dependent manner, consistent with previous studies (*19, 20*). Moreover, we benchmark the assay by directly comparing EVs and viruses, and resolving the fusion intermediates using cryogenic Transmission Electron Microscopy (Cryo-TEM) and Electron Tomography (Cryo-ET). We find that in EVs, contrary to most viruses, the low-pH-trigger is reversible, as previously shown for vesicular stomatitis virus, VSV (*22, 23*).

These results provide novel insight into the mechanism of EV fusion, suggesting that viral and EV fusogens likely share structural and functional similarities and may even share common ancestors. Moreover, they establish a standard method for further functional and structural studies of the fusion process that could lead to the identification of the EV fusion machinery and become a gold standard approach in the EV fusion field.

## Results

### EVs and large unilamellar vesicles (LUVs) can be distinguished using cryo-TEM

The delivery of the EV cargo into the host-cell cytoplasm must initiate with binding to the recipient cell membrane, followed by either fusion with the plasma membrane or vesicular uptake (i.e., endocytosis) of the EV, followed by fusion with the endosome (**Fig. 1 A**) (*12*). To bypass this complexity and focus on the fusion process, we probed the interaction of EVs with liposomes *in vitro*.

**Figure 1.**
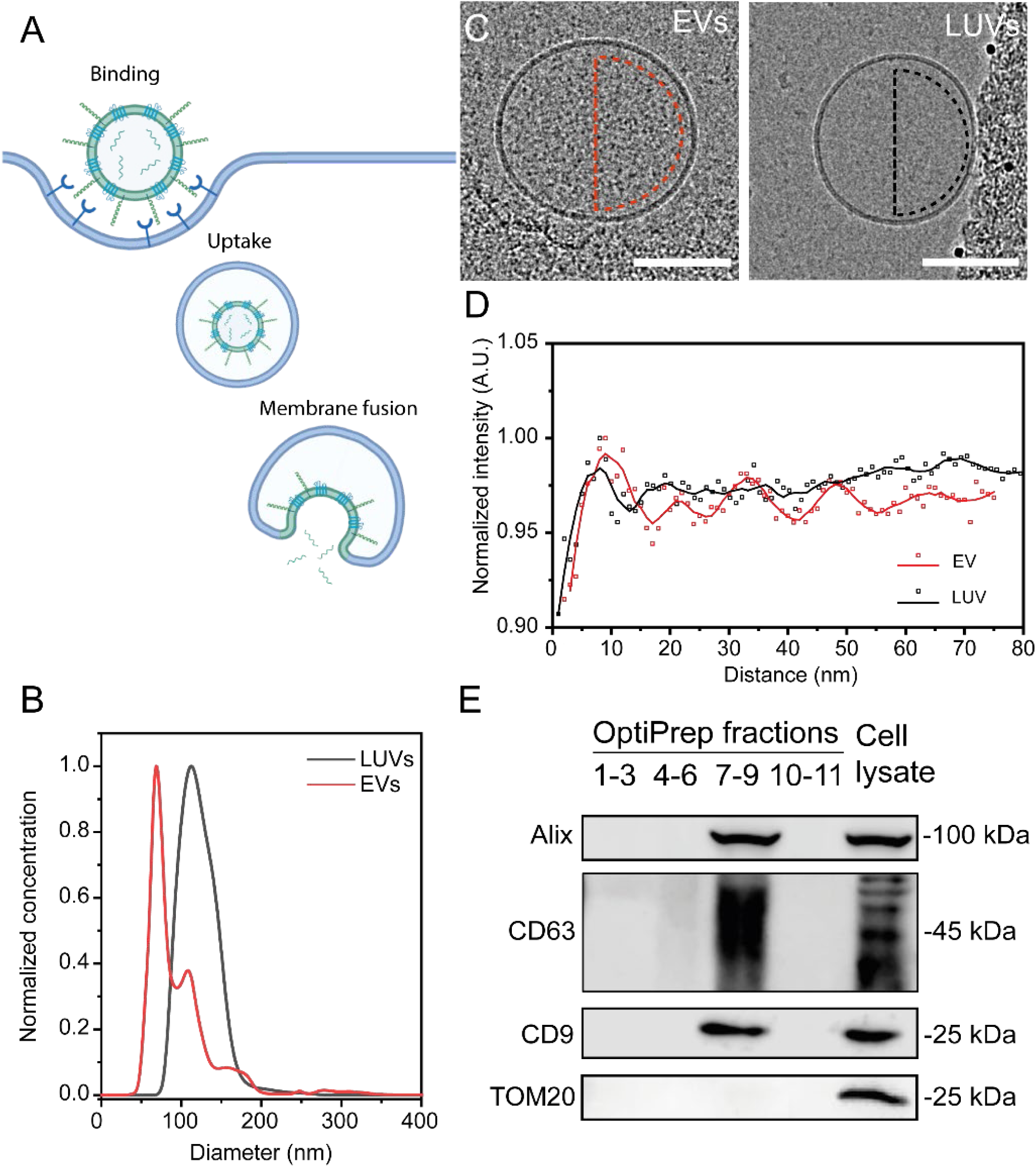
EV-LUV *in vitro* system to investigate membrane fusion. **(A)** Graphical illustration of the EV uptake/cargo delivery pathway. **(B)** Comparison of the size distribution of EVs (red) and LUVs (black) measured via NTA. **(C)** Representative Cryo-TEM image of EVs isolated from OVCAR-3 cell supernatants and LUVs. Scale bar 100nm. **(D)** Radial intensity profile of gray levels in the lumens of EVs (red) and LUVs (black). Dots represent data points obtained from radially averaged line profiles of gray level intensity. Lines represent data smoothed using a 25 points Savitzky-Golay filter. (**E**) Western blot analysis of density gradient fractions (pooled by density) and OVCAR-3 cell lysate (as a control) using antibodies against EV (Alix, CD9 and CD63) and control protein determinants (TOM20). Illustration created with BioRender.com.

We purified extracellular vesicles from OVCAR-3 cell culture supernatant using OptiPrep density gradient ultracentrifugation (*24*) following MISEV guidelines (*25*). We subsequently pooled the EV fractions and pelleted them through a sucrose cushion (*26*). We examined EV (and LUVs) samples size and morphology using nanoparticle tracking analysis (NTA) (**Fig 1 B and Figs. S1 A-B**) and Cryo – TEM (**Fig. 1 C**), showing that the isolated EVs display the typical morphology and size distribution of EVs with an average diameter of 132.5 ± 2.1 nm. We also detected the known EV markers CD63, Alix and CD9 (*25*), but not the mitochondria-specific protein TOM20, as expected with purified Evs (*25*) (**Fig. 1 E**). Isolation from naïve growth medium was used as a control to verify that EVs originate from the cultured cells and not from the bovine serum (**Fig. S1 B**).

LUVs extruded at 100 nm showed a narrow diameter distribution at 109.9 ± 2.6 nm (**Fig 1 C**). Importantly, we could distinguish EVs and LUVs in cryo-TEM by the spatial distribution of gray levels within each vesicle lumen (**Fig 1 C and D**). LUVs display a smooth distribution while EVs display a granular pattern characterized by periodicity in the radial intensity signal, consistent with the absence and presence of cargo, respectively (**Fig. 1 D**).

### Membrane mixing between EVs and LUVs is triggered by low pH

We then turned to methods extensively utilized in virology to study viral membrane fusion and established an *in vitro* membrane mixing essay based on FRET (**Fig. 2 A**) to probe EV membrane fusion with lipid membranes. (*27–32*). To benchmark the assay, we conducted the experiments with EVs and non-replicating retroviruses, stained with the lipid dyes DiI and DiD as a FRET pair. At steady-state, DiI fluorescence emission is transferred to and absorbed by DiD. The FRET-labelled vesicles or viruses were incubated with unlabeled LUVs. If fusion occurs, donor intensity increases due to the dilution of the vesicles or viruses by the unlabeled LUV membranes, which increases the distance between the FRET pair. The donor’s fluorescence (DiI) is monitored, and its intensity is normalized to the maximum donor intensity, which is obtained by fully solubilizing the membranes using a detergent (**Fig. 2 A**). To mimic the lipid composition of the late endosome membrane, we conducted this analysis using LUVs enriched in bisoleoyl-lysobisphosphatidic acid (LBPA) and without cholesterol (*33*). For controls, we used labeled retroviruses or LUVs mixed with unlabeled LUVs.

**Figure 2.**
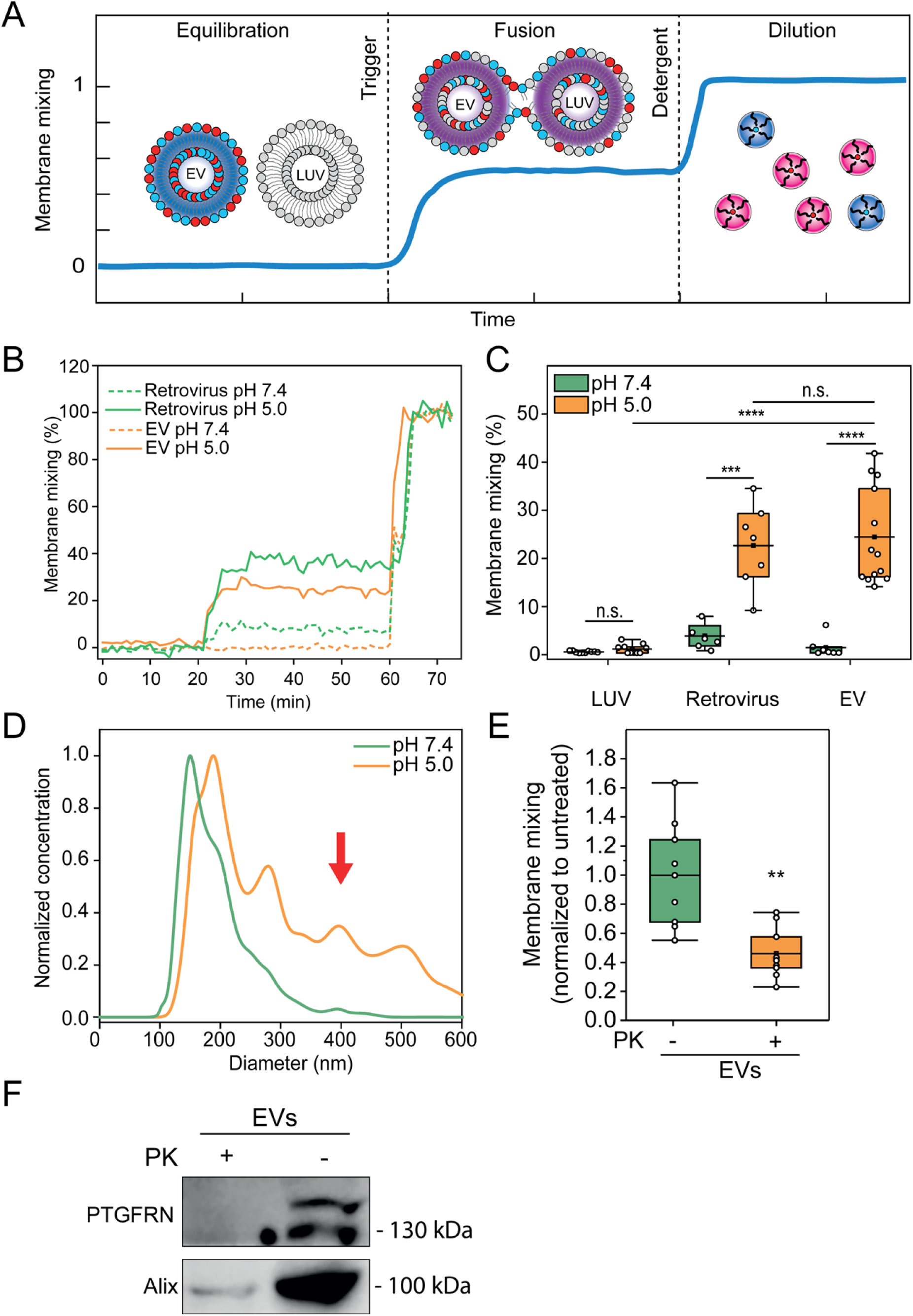
EV fuse in a protein and pH-dependent manner reminiscent of viruses. **(A)** Illustration of the FRET-based membrane mixing assay employed to quantify fusion between EVs and liposomes. Highly FRET-efficiency labeled EVs are incubated with non-labeled liposomes and their ability to fuse is probed by monitoring the donor fluorescent intensity after triggering. **(B)** Representative curves of membrane mixing assay for either retrovirus (green) or EV (orange) incubated with unlabeled LUVs at pH 7.4 (dotted line) or pH 5.0 (solid line). **(C)** FRET fusion assay for labeled retroviruses, EVs and LUVs, incubated with LUVs at either pH 7.4 (green) or pH 5.0 (orange), showing that retroviruses and EVs fuse with similar efficiencies. LUVs mimicked the late endosome lipid composition. **(D)** Representative NTA size distribution curves of EVs-LUVs mixtures upon incubation at pH 7.4 (green) or pH 5.0 (orange) showing increase in vesicle size after mixing and triggering with pH 5.0. **(E)** FRET membrane mixing assay comparing EVs treated with Proteinase K (PK, orange) or non-treated (green), incubated with late endosomal-mimicking LUVs at pH 5.0. **(F)** Western blot for membrane protein EV marker, PTGFRN, and intraluminal protein Alix for non-treated and PK – treated EVs, showing that proteinase only digests proteins on the surface of the EVs.

As pH acidification was shown to trigger viral (*23, 34*) and EV fusion (*19, 20*), we validated the assay by testing whether membrane mixing is pH-dependent. Incubation of the labeled EVs with unlabeled LUVs at pH 7.4 shows negligible membrane mixing, indicating no fusion occurs (**Fig. 2 B**). Upon acidification (ranging from pH 7.4 to 5.0), we observed significant membrane mixing for EVs and viruses at comparable efficiencies (p = 0.699; 24.4 ± 10.0 % and 22.7 ± 8.7 % respectively; **Fig. 2 C and Figs. S2 A**). LUV control showed no significant variation in membrane mixing across the range of acidic pH values, indicating that pH alone is not sufficient to induce fusion and lipid mixing (**Fig. S2 A**). Additionally, no membrane mixing is measured in fractions and naïve growth medium absent of EVs (**Fig. S2 B-C**).

If fusion occurs between EVs and LUVs, then the size distribution of the population is expected to skew towards larger vesicles upon acidification. To test size distributions under different pH conditions directly, we performed NTA analysis on the mixed EVs and LUVs at pH 7.4 compared to pH 5.0. We observed that at pH 7.4, the diameters are consistent with a mixed population of EVs and LUVs. However, the diameters shift towards larger sizes after acidification, consistent with vesicle fusion (219 ± 20 nm at pH 7.4, 300 ± 30 nm at pH 5.0, p = 2,36E-9; **Fig. 2 D and Fig. S2 D**). Together, these results demonstrate that EVs fuse in a process triggered by low pH, similar to viruses.

### EV proteins and target membrane lipid composition are essential for fusion

Next, we examined if EV fusion is protein mediated by proteolytically “shaving” the proteins from the EV membrane using Proteinase K (PK), a broad-spectrum serine protease (*35, 36*). We observed a significant reduction in membrane mixing of EVs shaved with PK (46 ± 18 % compared to NT EVs; p = 0.00104 **Fig. 2 E**). PK treatment showed no significant vesicle size distribution or concentration alteration as measured by NTA (**Fig. S2 E**). Fusion efficiency was also not affected by treatment with the protease inhibitor phenylmethylsulfonyl fluoride (PMSF), which was used to quench PK digestion (**Fig. S2 F**). These results suggest that EV membrane proteins are critical for the fusion mechanism, consistent with previous reports showing that EV content release is protein-dependent (*19*). We next verified by western blot that PK digestion removed the EV membrane protein PTGFRN (*37*) but retained the luminal protein Alix, showing that EV integrity was maintained and only surface proteins were digested (**Fig. 2 F**).

EVs fuse to late endosomal-mimicking membranes at pH 5.0 (**Fig. 2 C**) with an efficiency comparable to viruses. To investigate whether the late-endosomal lipid LBPA is essential for efficient fusion, we examined whether EVs could fuse at a similar probability to either single-component lipid bilayer (DOPC) or early endosomal-mimicking membranes. We found that fusion does not occur when EVs interact with non-physiological DOPC LUVs (**Fig. 3 A**, 1.0 ± 1.1 %), reinforcing the concept that lipid composition is crucial for membrane fusion (*38*). Moreover, we observed significantly lower fusion efficiency with early endosomal-mimicking LUVs compared to late endosome composition (9.2 ± 5.3 % and 28.2 ± 10.1 % respectively; p = 0.000125). This effect may arise from cholesterol in the bilayer, which was reported to inhibit efficient cargo transfer from EVs to recipient cells at the late endosome (*17*).

**Figure 3.**
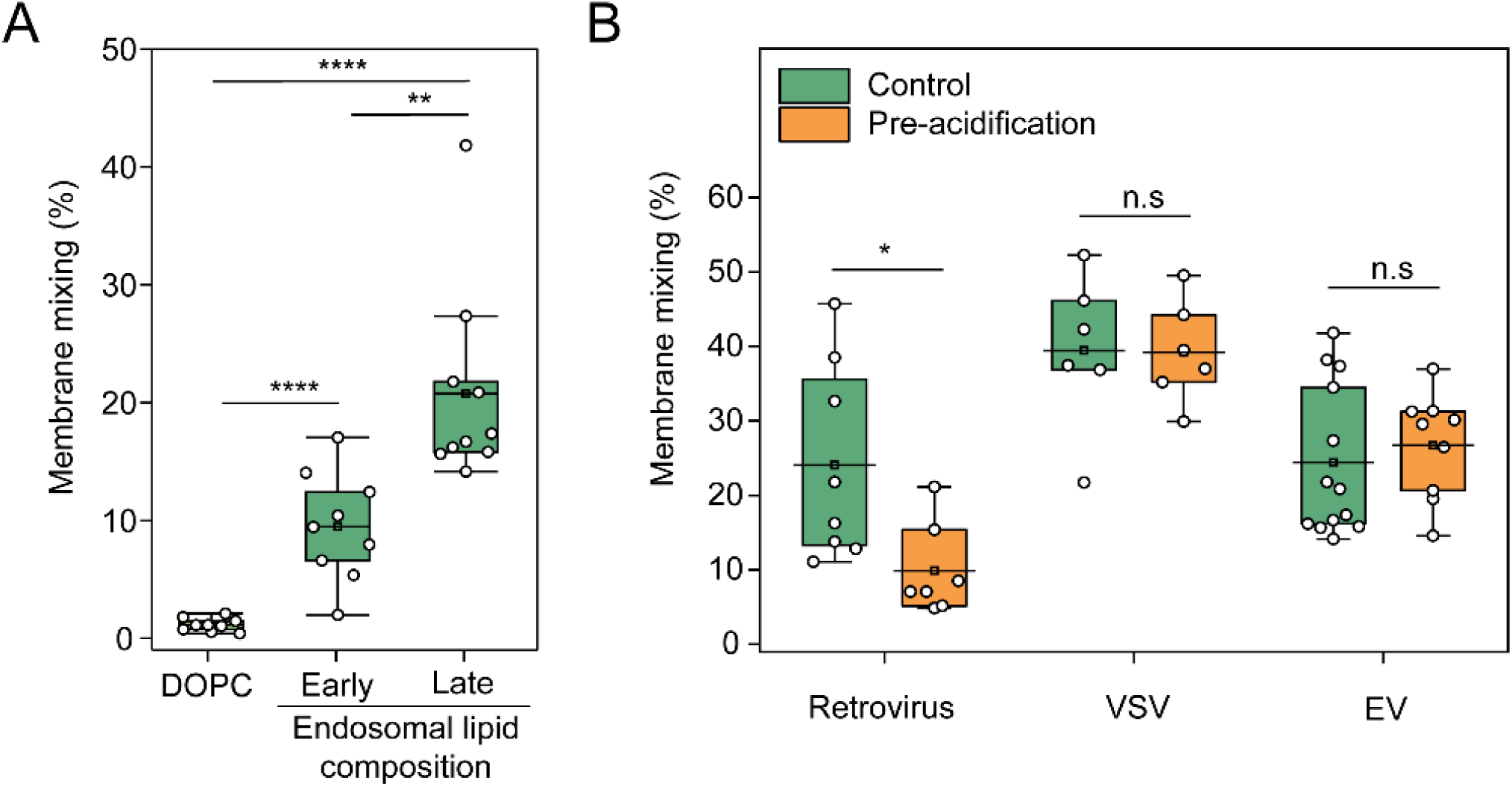
EV fuse using a lipid-composition dependent but reversible mechanism. **(A)** FRET fusion assay for EVs incubated with LUVs composed of DOPC (DOPC), early endosome-mimicking LUVs, or late endosome-mimicking LUVs at pH 5.0. **(B)** Membrane mixing assay to probe reversibility of the putative EV fusogen. EVs, Retroviruses, and VSV viruses were incubated at pH 5.0 and subsequently brought back to pH 7.4 and mixed with LUVs at pH 5.0.

### The putative EV fusogen is insensitive to pre-acidification

Viral fusogens typically undergo an irreversible conformational change at low pH (*39*). Hence, exposing retroviruses to low pH in the absence of target membranes inactivates the fusogen. Since the biogenesis of specific EV subpopulations occurs in the acidic milieu of MVBs (e.g., exosomes) (*40*), we hypothesized that the conformational change of the putative EV fusogen might be reversible, as shown for the viral fusogen VSV-G (*22, 41, 42*). To test this hypothesis, we measured the ability of EVs to fuse after acidification and subsequent re-neutralization of the pH values prior to incubation with LUVs and reacidification. Membrane mixing efficiency was evaluated under these conditions and compared to both pseudotyped VSV and retroviruses as positive and negative controls, respectively (*43*). While retroviruses lost their membrane mixing activity after acidification and re-neutralization, VSV and EVs exhibited comparable membrane mixing probabilities in the two conditions (**Fig. 3 B and S3**). We conclude that the putative EV fusogen is triggered by low pH but in a reversible manner. These results are consistent with a model wherein EVs are not fusogenic during their biogenesis in the acidic lumen of the MVB, and only become primed for fusion upon release into the neutral pH of the extracellular space. Moreover, they imply that the glycoproteins on the EV surface may be structurally similar to VSV-G.

### EV fusion intermediates visualized by Cryo-TEM

Having demonstrated that the *in vitro* EV-LUV system recapitulates the previously reported protein- and pH-dependence, we used Cryo-TEM to visualize EV - LUV interactions (**Fig. 4**). Membrane fusion intermediate states canonically associated with viral fusion include (i) close contact between the lipid bilayers, (ii) fusion of the outer leaflets to form a hemifusion diaphragm, (iii) fusion of the inner leaflets to allow content mixing, and (iv) expansion of the fusion pore (*44–46*) (**Fig. 4 A**). EV incubated with LUV at pH 5.0 and 7.4 showed a similar percentage of close contacts between the two vesicle populations (38.5 ± 16.6 % and 26.2 ± 8.6 % respectively; **Fig. 4 B and F**). Remarkably, fusion intermediates including hemifusion (**Fig. 4 C and F**; 6.8 ± 2.8 %), content mixing (**Fig. 4 D and F**; 10.4 ± 5.9 %) and expanded pore (**Fig. 4 E and F**; 8.8 ± 1.6 %) were only apparent at pH 5.0. LUVs alone displayed some close contacts but no fusion intermediates at both pH 7.4 and 5.0 (6.9 ± 2.6 % and 5.4 ± 0.9 % respectively; **Fig. 4 F and Fig. S4 A**).

**Figure 4.**
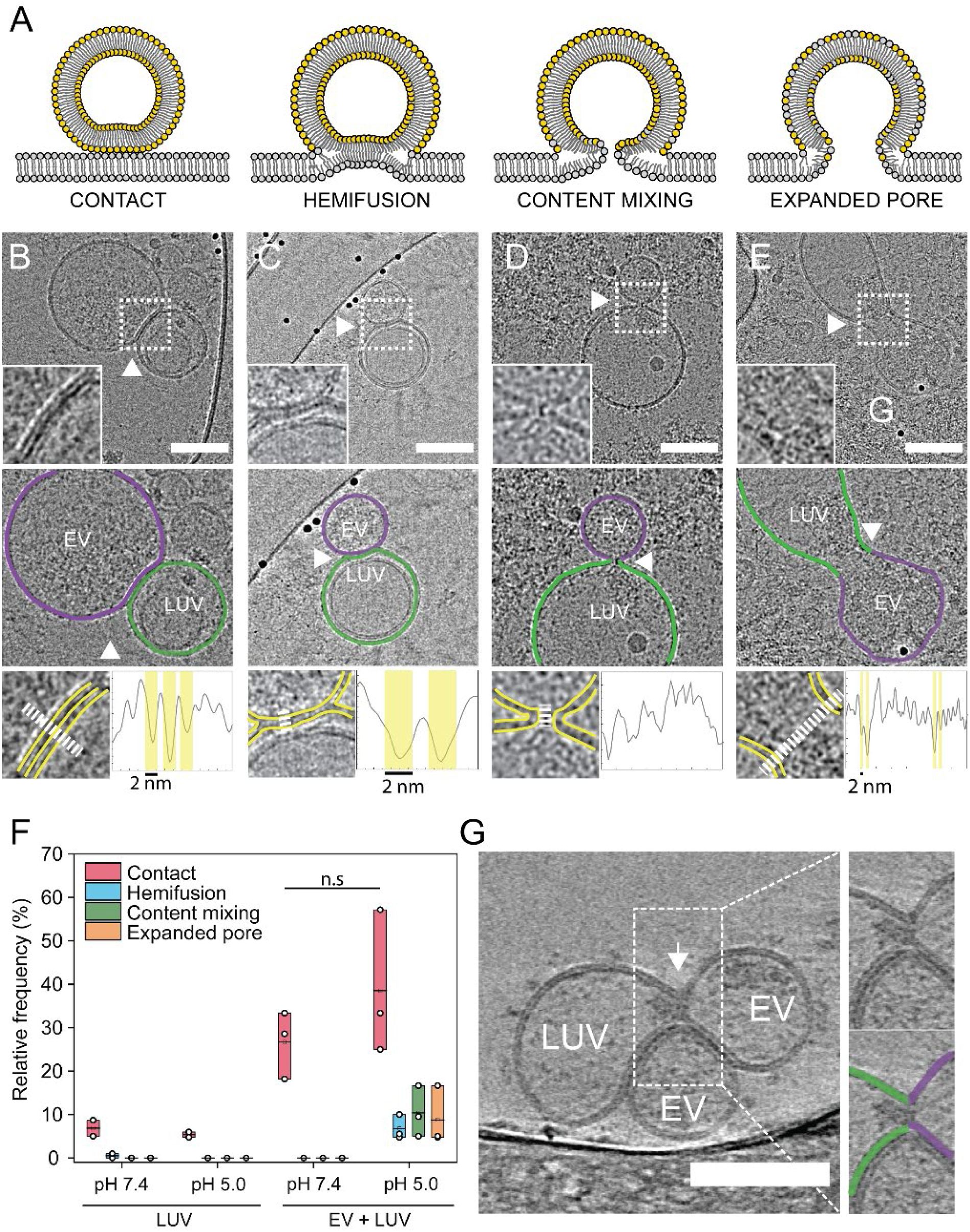
Cryo-EM imaging reveals EV hemifusion intermediates. **(A)** Illustration showing canonical membrane fusion intermediates. Contact: the viral membrane is tightly in contact (< 2 nm) with the apposing membrane and the two bilayers run parallel to each other. Hemifusion: the two proximal leaflets of lipid bilayers have joined and the hemifusion diaphragm is composed of only the two remaining leaflets. Content mixing: the two membranes merge at the contact point with the two bilayers transitioning continuously from one onto the other. At this stage, the content can mix, but the pores can still collapse and reseal. Expanded pore: the fusion pore increases in diameter and complete content mixing can occur. **(B – E)** Representative Cryo – TEM images of EV-LUV fusion intermediates. Insets show the interaction spot at higher magnification (Top panel). Middle panel: Outlines showing EV (magenta) and LUV (green) membranes as defined by luminal gray level distribution. Bottom panel: Line profile to evaluate the presence of bilayer-leaflets. Yellow and white lines indicate membrane leaflets in inset and region where line profile was acquired, respectively. **(B)** Contact between EV and LUV **(C)** Hemifusion. Arrow indicates the location where the two bilayers merged into one. **(D)** Content mixing. White arrow indicates the fusion pore. **(E)** Expanded pore: dumbbell-shaped pore with an enlarged neck and apparent flow of the EV cargo into the LUV lumen. White arrow indicates the putative extended neck where the vesicles fused. **(F)** Quantification of interaction intermediates for LUVs – LUVs and EVs – LUVs systems at pH 7.4 and 5.0. **(G)** Tomographic reconstruction of EVs (magenta) – LUV (green) interaction at pH 5.0 revealing content mixing between the vesicles. White arrowhead indicates the fusion pore.

To resolve the 3D ultrastructure of the fusion intermediates and unambiguously determine if content mixing occurs through an expanded pore, we used Cryo-ET. Reconstructed tilt series of acidified EVs with LUVs clearly showed content mixing and expanded pore between EVs and LUVs, with the two membranes fully merged and a narrow connection between the two vesicular lumens (**Fig. 4 G, S4 B and Movie S1, S2**). These results demonstrate that EV fusion is triggered by low pH and that membrane mixing using FRET is a *bona fide* method to measure fusion and the efficiency of cargo delivery under varying conditions.

## Discussion

While it has not been unambiguously shown, the hypothesis that EV cargo delivery occurs via membrane fusion is supported by several studies (*17–20*). Yao *et al*. showed that membrane mixing of labeled exosomes occurs at the endosomes and that mixing depends on the late endosomal lipid LBPA (*18*). Similarly, Joshi *et al*. demonstrated that release of GFP from the EV lumen occurs at the late endosome and that inhibiting endosome acidification or cholesterol depletion suppresses EV cargo delivery (*17*). In two separate studies, Bonsergent *et al*. demonstrated *in vitro* that EV content release to plasma membrane sheets is protein and pH-dependent and that the delivery can be inhibited by IFITM proteins on the apposing membrane (*19, 20*). However, these assays could not unambiguously demonstrate that membrane fusion between EV and membranes occurs.

To overcome this challenge, we standardized a robust *in vitro* fusion assay between EV and LUVs and benchmarked it using viruses. We show that the assay recapitulates all the features previously observed for EV cargo delivery and allows better control over both the environmental conditions and the lipid composition of the target membrane. Moreover, the assay and fusion process can be readily visualized by Cryo-EM and Cryo-ET.

Taking advantage of these advantages, we show that EV fusion is unidirectional *in vitro* and does not require proteins on the target membrane. Moreover, fusion occurs via hemifusion. These features are strikingly similar to viral fusion, suggesting that the EV fusogen or fusogens might share structural similarities, or even common ancestors, with viral fusogens. Moreover, we demonstrate that changes in the delivery efficiency reported for specific lipid compositions (*17*) or upon pH acidification (*19*) are directly related to EV fusion.

Remarkably, our results also show that EVs are not inactivated if exposed to acidic pH in the absence of target membranes, suggesting a reversible conformational change of the EV fusogen, similarly to the viral fusogen VSV-G. These results are consistent with a model wherein the EVs (e.g., exosomes) that bud into the acidic lumen of the MVB are in an inactive non-fusogenic state. EVs are then activated at neutral pH after secretion to the extracellular space and fuse in a pH-dependent manner after internalization into target cells. Thus, EVs avoid a paradoxical scenario wherein exposure to acidic pH during assembly would also irreversibly inactivate their fusion machinery.

We have previously suggested that EVs derived from malaria-infected red blood cells (RBCs) can fuse to LUVs that mimic the lipid composition of the plasma membrane in a pH-independent manner (*47*). The absence of a pH trigger is consistent with the lack of endocytosis in RBCs, and suggests that the fusion mechanism and the fusogens might be context-dependent. This hypothesis further reinforces the analogy to viruses that have evolved different triggering mechanisms. Therefore, it will be essential to define the fusion mechanism of different subpopulations of EVs and identify the EV fusogens. However, better separation of EVs to structurally and functionally distinct subpopulations remains a confounding challenge.

A benchmarked assay that recapitulates *bone fide* EV fusion could become an essential tool in identifying the fusion machinery. Candidate proteins could be deleted in producing cells and EV fusogenicity could be evaluated with high throughput. Moreover, showing that isolated EVs maintain their fusogenic activity has the potential to become a gold standard in the study of EVs (*25*). Understanding the mechanism of EV membrane fusion is essential not only for expanding our knowledge in EV biology but also for developing them into biocompatible and tissue-specific delivery systems.

## Materials and Methods

### Statistical analysis

All experiments were carried out with n≥3 biological replicates. Statistical analysis was carried out using OriginPro software. In all Figures containing box plots, each dot represents one measurement. Box layouts represent 25 – 75 percentiles of the distribution, whiskers highlight outliers data points, and horizontal black lines represent mean of the distribution. Whenever comparing two conditions, data were analyzed with a two-sample student’s t-test with a significance level of 0.05. Throughout the study, the threshold for statistical significance was considered for p-values ≤0.05, denoted by one asterisk (*), two (**) if p ≤0.01, three (***) if p <0.001 and four (****) if p ≤0.0001.

### Cell culture and EV isolation

Extracellular vesicles derived from ovarian cancer cells (OVCAR-3; ATCC HTB - 161) conditioned media were harvested as previously described (*48*). Briefly, cells were seeded at 10×10^6^ cells in a 175 cm^2^ flask in culture media composed of DMEM with 10% EV-free FBS, 1% Sodium Pyruvate, 1% L-Glutamate and 1% PenStrep. When cells had reached 70% confluence (typically two days post-seeding), the cells were washed twice with PBS −/− and replenished with naive EV-free growth medium. Cell culture media was collected after 48 hours and spun at 300g for 10 min at 4°C to remove large debris and leftover cells, supernatant was collected and spun at 2,000g for 10 min at 4°C. Supernatant was then collected, spun at 10,000g for 45 min to remove larger vesicular particles, and filtered through a 0.22 µm polycarbonate filter. The resulting media was used for vesicles isolation within 2 days or frozen at −80°C for biochemical analysis.

Following filtration, the cell culture media was spun using an ultracentrifuge, a Ti45 rotor (Beckman Coulter, Fullerton, CA, USA) at 100,000 g for 4 h at 4°C. Supernatant was removed and the resulting pellet was washed once with PBS −/− and resuspended in PBS −/−. For membrane mixing experiments, cell culture media was incubated with 0.01% v/v of 2.5 mM DiI (Merck, CAT: 42364) and DiD (Thermo Fisher, D7757)) 1:1 mixture in DMSO at 37°C for 30 min.

### Density gradient ultracentrifugation

Following differential ultracentrifugation, EVs were fractionated by OptiPrep density gradient ultracentrifugation (100,000 × g, 18 h, 4 °C) using a SW41 rotor (Beckman Coulter, Fullerton, CA, USA) through a continuous 5–40% OptiPrep (Sigma-Aldrich, D1556) gradient. Fractions (1 ml) were collected from the top of the gradient for further analysis, and density was verified by measuring the mass of a 100 µL aliquot of each fraction. Fractions of EV-specific density were then pooled together and subsequently concentrated via ultracentrifugation (100,000 × g, 4 h, 4 °C) through a 20% w/v sucrose cushion in a SW41 rotor (Beckman Coulter, Fullerton, CA, USA). The resulting supernatant was discarded, and the EV pellet was resuspended in PBS −/−.

### Preparation of vesicle-depleted Fetal Bovine Serum (EV-free FBS)

FBS was depleted from extracellular vesicles by two rounds of ultra-centrifugation at 100,000 g for 18 h in a Beckman Ti45 rotor; each round the supernatant was collected and the large pellet at the bottom of the tube discarded. After the final round of ultracentrifugation, supernatant was collected and filtered through a 0.22 µm pore membrane, aliquoted and stored at −20°C for preparation of EV-free growth medium.

### Nanoparticle tracking analysis

EV size and concentration distribution analysis was performed using nanoparticle tracking analysis (Malvern Instruments Ltd., NanoSight NS300) at 20 °C. Sample size distributions were obtained in a liquid suspension (1:500 - 1:1000 dilution in PBS −/−) by analyzing Brownian motion via light scattering. The camera level was set to 13 and gain to 1, with a 405 nm laser unit without filter, following the manufacturer’s instruction. The data was analyzed using NTA 2·1 software (NanoSight) and plotted using the OriginPro software.

### Western blot analysis

Equal volumes of pelleted OptiPrep fractions and 20-30 µg of protein cell lysates were mixed with 4x Laemmli sample buffer (4% SDS, 10% mercaptoethanol, 20% glycerol, 0.004% bromophenol blue and 0.125M Tris-HCl) and boiled at 96°C for 5-10 minutes. Samples were subjected to electrophoresis using 7-15% SDS-PAGE gels in TG-SDS running buffer (Bio-Lab) at constant 150 V for 1 h. Proteins were electrotransferred onto nitrocellulose membranes using a standard tank transfer protocol with TG transfer buffer (Bio-Lab) with 20% methanol. Membranes were blocked with 5% non-fat milk dissolved in TBS containing 0.1% Tween (TBST) for one hour and incubated with one of the following primary antibodies either overnight at 4°C or 1 hour at room temperature. (dilution, company, catalog number): anti-CD9 (1:1000, Abcam, ab92726), anti-PTGFRN (1:1000, R&D Systems, MAB10043-100), anti-CD63 (1:1000, Proteintech, 25682-1-AP), anti-Alix (1:1 000, Proteintech, 12422-1-AP), anti-Tom20 (1:1 000, Abcam, ab56783). The primary antibodies were diluted in 5% non-fat dry milk in TBST. Membranes were washed 3 times for 10 minutes at room temperature with TBST and incubated with either anti-Mouse IgG-HRP (1:20 000, Abcam, ab6728) or anti-Rabbit IgG-HRP (1:20 000, Abcam, ab6721) diluted in TBST for 1 hour. Membranes were washed 3 times for 10 minutes with TBST. EZ-ECL (Biological Industries Ltd.) was used for detection with the sequential visualization using the Odyssey Fc Dual-Mode Imaging System (Li-COR Biosciences, USA). Each presented western blot is a representative image of three separate biological replicates.

### Preparation of large unilamellar vesicles (LUVs)

LUVs were prepared with a lipid composition either of DOPC or mimicking the membranes of early and late endosomes. For early endosome-mimicking LUVs, the lipid content was DOPC:DOPE:SM:chol 30:10:25:35, while late endosome-mimicking LUVs were composed of LBPA:DOPE:DOPC 70:5:25 (molar ratio) to mimic the cholesterol sequestration and enrichment of late endosomal lipid LBPA (*49*). Lipid solutions in chloroform of the different phospholipid species were mixed to the desired molar ratios in a glass vial, and the organic solvent was evaporated by 12 h of vacuum pumping. For labeled LUVs, the lipids were stained at a 2 % mol/mol fraction of DiI and DiD in chloroform before evaporation. The lipid film was then hydrated with PBS −/− at 50 °C to reach the desired concentration and gently vortexed. The resulting MLV suspension was then sonicated for 10 min to disperse larger aggregates and the liposomal suspension was extruded 21 times through polycarbonate filters (100 nm pore size, Avanti Polar Lipids) using a mini-extruder (Avanti Polar Lipids). Size and concentration were verified using NTA and the liposomal suspension was used within 2 weeks from extrusion.

### Retroviruses preparation

Retroviruses were generated by transfecting pBABE-Puro plasmids, a gift from Hartmut Land & Jay Morgenstern & Bob Weinberg (Addgene plasmid # 1764, (*50*)), into Platinum-E Cells (Cell Biolabs, Inc.). 24hrs prior to transfection, 3× 10^6^ were seeded in 10cm culture dish according to manufacturer instructions. 10μg of retroviral plasmid DNA was transfected using jetPRIME® transfection reagent (PolyPlus transfection). 5ml of viral suspension was collected from the conditioned media 48hrs post-transfection and centrifuged at 1,000 g for 10 min at 4°C to remove cell debris. The supernatant was carefully transferred into another ice-cooled falcon tube. Virions were concentrated by pelleting at 100,000 g through a 20% sucrose cushion for 2 h and resuspended in PBS −/− The concentrated viruses were used for further experiments.

### Preparation of VSVΔG-G pseudoviruses

Baby Hamster Kidney cells (BHK-21; ATCC) were grown in Dulbecco’s modified Eagle’s medium (DMEM, Gibco), 1% Penn/Strep, 7-10% Fetal Bovine Serum (FBS, Biological Industries, Kibbutz Beit Haemek, Israel) at 37°C in 5% CO_2_. For maintenance, BHK-21 cells were grown at 7% FBS. For pseudovirus preparations, BHK-21 cells were grown at 10% FBS.

To generate VSV-G-complemented VSVΔG pseudoviruses (VSVΔG-G), 200,000 BHK-21 cells were seeded in 5 ml of medium. Cells were transfected at ∼70% confluency with plasmids encoding VSV-G (Indiana) glycoprotein (1 μg/ml) (*51*). After 24 hours incubation, transfected cells were infected with VSVΔG-G helper viruses at a multiplicity of infection (MOI) of 5 for 1 hour at 37°C in a 5% CO_2_ incubator rocking every 15 minutes. After 1 hr, the medium was replaced with serum-free medium. 24 hours post-infection, the cells were scraped off and transferred with the supernatant into ice-cooled falcon tubes. The cell debris were removed by centrifuge at 500 g for 10 mins at 4 °C. The supernatant was carefully transferred into another ice-cooled falcon tube. Virions were concentrated by pelleting at 100,000 g through a 20% sucrose cushion for 2 h and resuspended in PBS −/− The concentrated viruses were used for further experiments.

### Pre-acidification of virions and EVs

EVs or viruses’ samples post-isolation were acidified to pH 5.0 by adding 7% v/v of HCl 100 mM and incubated at 4°C for 45 min. Subsequently, 6.5% v/v of NaOH 100 mM was added to re-equilibrate the pH to 7.4 and samples were maintained at 4°C for at least 1 hour before being incubated with liposomes for membrane mixing assay (*39, 52*).

### Membrane mixing assay

All experiments were performed using a Cytation 5 Imaging Reader plate reader (BioTek) with a 96-well plate. DiI (Merck, CAT: 42364) and DiD (Thermo Fisher, D7757) labeled EVs and unlabeled LUVs were diluted in PBS −/− per well to reach a final ratio of 1:9 fluorescent particles to non-labeled vesicles, and fluorescence intensity of the donor (DiI) was recorded every 60 s for 30 min, with excitation wavelength of 530 nm and emission wavelengths of 570 nm. Subsequently, a volume fraction of HCl 100mM was added to reach the desired pH and DiI fluorescence intensity was recorded for 1 h every 60 s. Finally, Triton X-100 was added in each well to reach 0.1% final concentration and fluorescence intensity was recorded for 15 min every 60 s. The emission fluorescence for each time point was measured as I_n_. The emission fluorescence of the untreated liposomes was measured as I0, and that of the liposomes solubilized with 0.1% TRITON X-100 was defined as I_100_. The percentage of membrane mixing at each time point is defined as: donor relative intensity (% of TRITON X-100) = (I_n_ − I_0_) × 100/(I_100_ − I_0_). All measurements were performed at 37 °C. The data was analyzed by using Gen5™ v. 3.04 software (BioTek).

### Size distribution NTA analysis for EV fusion events

LUVs and EVs were mixed in a 1:1 ratio (particles:particles) to a final concentration of 1–10 × 10^8^ particles/mL in 1mL of filtered PBS −/− and kept at 4°C for 1 hour. Prior to size NTA measurement, a 7% v/v fraction of either PBS −/− or HCl 100 mM was added to maintain physiological pH or reach pH 5.0, respectively. Samples were incubated at 37°C for 30 min and the size distribution was subsequently measured using a NanoSight NS300. Briefly, approximately 1ml solution was loaded into the sample chamber of an LM10 unit (NanoSight) and five videos of 60 s were recorded. Data analysis was performed with NTA 2·1 software (NanoSight). The resulting size distribution curves were then analyzed by considering the average diameter of each biological repeat, obtained from the instrumentation analysis software.

### EV protein digestion by Proteinase K

Isolated EVs were incubated for 45 min at 37°C in the presence of 20 μg/ml Proteinase K (Invitrogen, AM2546). Following incubation, the sample was placed on ice and the proteinase activity was quenched with 2 mM phenylmethylsulfonyl fluoride (Merck, P7626) in DMSO.

### Preparation of Cryo-TEM samples

Cryo-EM samples of both EVs and LUVs were prepared on either lacey carbon or C – flat EM grids (Electron Microscopy Sciences, USA), on which 10 nm Protein A colloidal gold particles (Au – NP) were pre-adsorbed (Aurion, Netherlands). Au – NP adsorbed grids were then glow-discharged (30 s, 25 mA) in a Pelco EasiGlow system. An aliquot (3.5 μL) of the aqueous solution of the sample was applied on to the carbon side of EM grids, which was then incubated in the humidity chamber of the instrument for 7 min at 100% humidity and room temperature, and subsequently blotted for 4.0 s at blot force −10 and plunge-frozen into the precooled liquid ethane with a Vitrobot Mark IV (FEI, USA).

### Cryo-TEM

Cryo-electron micrographs of vitrified samples were collected using a transmission electron microscope Talos Arctica G3 TEM/STEM (Thermo Fisher Scientific, USA), equipped with a OneView camera (Gatan) at accelerating voltage of 200 kV. Grid mapping and image acquisition were performed using SerialEM software (*53*) at a nominal magnification of x180 and x13,500, respectively. High magnification images were recorded at x73,000 nominal magnification (0.411 nm pixel size) with a −3.5 µm defocus value. To minimize radiation damage during image acquisition, low-dose mode in SerialEM software was used and electron dose was kept below 100 e^−^ Å^−2^.

### Cryo-electron tomography (cryo-ET)

Samples were prepared as for cryo-TEM (described above) with some modifications. Prior to plunging, samples were mixed 50:1 with a suspension of 10-nm Au – NP (Aurion, Netherlands) to serve as fiducial markers for reconstruction. Tilt series were collected using a transmission electron microscope Titan Krios 3Gi STEM/TEM microscope (Thermo Fisher Scientific, USA) at 300 kV equipped with a Gatan K3 direct detector mounted at the end of a Gatan BioQuantum energy filter set in zero-energy-loss mode (slit width, 20 eV). Tilt series were acquired in low dose mode using SerialEM (*53*) software at a nominal magnification of ×42,000 with an angular range from −60° to +60°, an angular increment of 4° using a −3.5 µm defocus, 70 µm objective aperture, 0.214 nm per pixel and a maximal total dose of 150 e^−^ Å^−2^. Tomograms were reconstructed using the weighted back-projection technique in the IMOD software suite (*54*) with a SIRT-like filter equivalent to 5 iterations, following nonlinear anisotropic diffusion (NAD) de-noising (*55*) if indicated.

### Cryo-EM image analysis of vesicles

The spatial distribution of gray levels of vesicles’ luminal content was analyzed from images collected both via Cryo-TEM or reconstructed Cryo-ET tilt series using the Radial Profile Angle plugin of Fiji (*56*). The obtained gray level intensity profiles, averaged over the radial angle of the lumen, were subsequently smoothed using a 25 points Savitzky-Golay filter on the OriginPro software to obtain the trend of the spatial distribution.

## Supporting information

Original Gels of Western Blot figures

## Acknowledgments

We thank Benjamin Podbilewicz for reagents and members of the Avinoam, Regev-Rudzki, and Gil labs for discussions. This research was supported by the ISRAEL SCIENCE FOUNDATION (grant No. 1637/20), within the Israel Precision Medicine Partnership (IPMP) program to O.A., N.R.R and Z.G. This study was also supported by the European Research Council (ERC StG # 851080 to O.A.). O.A. also acknowledges funding from the David Barton Center for Research on the Chemistry of Life and the Ruth and Herman Albert Scholarship Program for New Scientists. O.A. is an incumbent of the Miriam Berman Presidential Development Chair. The research of N.R.R. is supported by the Benoziyo Endowment Fund for the Advancement of Science, the Jeanne and Joseph Nissim Foundation for Life Sciences Research and the Samuel M. Soref and Helene K. Soref Foundation. N.R.R is the incumbent of the Enid Barden and Aaron J. Jade President’s Development Chair for New Scientists in Memory of Cantor John Y. Jade. N.R.R is grateful for the support from the European Research Council (ERC) (grant agreement No. 757743), the Weizmann - Sao Paulo Research Foundation (FAPESP) Brazil; supported by a research grant from the Instituto Serrapilheira and the Israel Science Foundation (ISF) (Grant Application no. 570/21). M.I.M was supported by the Lombroso Postdoctoral Fellowship for Cancer Research.

## Author Contributions

M.I.M, N.R.R and O.A. conceived and designed the experiments. M.I.M and P.B performed most of the experiments. E.O.P, S.K, G.Z, P.A.B and Y.E.A produced viruses and assisted in cell maintenance and biochemistry. T.N, L. G, Z.G. assisted with EV assays. M.I.M, N.R.R and O.A. wrote the manuscript with inputs from all authors.

