## Supplementary Information for "Extracellular vesicle fusion visualized by cryo-EM"

**This PDF file includes:**

Figs. S1 to S4
Captions for Movies S1 to S2

**Other Supplementary Materials for this manuscript include the following:**

Movies S1 to S2

<u>**SUPPLEMENTARY FIGURES**</u>

**Figure S1. (A)** EVs isolated via ultracentrifugation (green) compared to density gradient fractions grouped by density: fractions 1 - 3 (yellow) corresponding to 5% OptiPrep, fractions 4 - 6 (blue) corresponding to 10% OptiPrep, fractions 7 - 9 (orange) corresponding to 20% OptiPrep, and fractions 10 - 11 (gray) corresponding to 40% OptiPrep. Fractions 7 – 9 display a similar size distribution and concentration of particles to EVs isolated by ultracentrifugation. **(B)** Comparison of NTA size distribution of isolated sample from cell culture supernatant (green) and naïve growth media (orange) after differential ultracentrifugation.

**Figure S2. (A)** FRET fusion assay results for labeled retroviruses (green), EVs (orange), and LUVs (gray) incubated with unlabeled LUVs at pH ranging from 7.4 to 5.0, showing that a significant increase of the donor intensity occurs only upon progressive acidification. **(B)** FRET fusion assay for EVs isolated from pooled density gradient fractions and via ultracentrifugation only, incubated with LUVs at pH 5.0. Only fractions 7 – 9 show a similar membrane mixing compared to control EVs. **(C)** FRET fusion assay for OVCAR-3 cell culture media and naïve growth media, after ultracentrifugation-based isolation protocol (see Materials and Methods), incubated with late endosomal LUVs at pH 7.4 (green) and 5.0 (orange), showing that a significant increase of the donor intensity occurs only for samples isolated from cell supernatant, but not from growth media. **(D)** Mean diameter of suspension of either LUVs only or EVs incubated with LUVs, at pH 7.4 (green) or pH 5.0 (orange). **(E)** Representative NTA size distribution curve comparing non-treated EVs (green) and EVs treated with Proteinase K at 37°C for 45 min (orange). **(F)** FRET fusion assay comparing EVs either treated with proteinase inhibitor PMSF or non-treated, incubated with late endosomal-mimicking LUVs at pH 5.0. In all panels, late endosomal-mimicking LUVs were utilized.

**Figure S3.** Representative NTA size distribution curves comparing non-treated EVs (green) and EVs pre-acidified at pH 5.0 without incubation with LUVs for 45 min at 4 °C (orange).

**Figure S4. (A)** Representative Cryo-EM images of LUVs at pH 7.4 (top) and pH 5.0 (bottom). At pH 7.4 LUVs do not significantly interact and display only minor contact sites with no extended interface between bilayers. Upon acidification, no fusion intermediates are observed. Scale bars 100 nm. **(B)** Tomographic reconstruction of EV (magenta) – LUV (green) interaction at pH 5.0 reveals an expanded pore intermediate, with the EV cargo leaking into the liposomal lumen. Scale bar 100 nm.

**Movie S1.**

Tomogram showing content mixing between two EVs and a single LUV at pH 5.0, corresponding to Fig. 4 G. Tomogram was acquired at defocus –3.5 and denoised using NAD filter with k value 5 and 5 iterations. Scale bar: 100 nm.

**Movie S2.**

Tomogram showing an expanded pore between EV and LUV at pH 5.0, corresponding to Fig. S4 B. Tomogram was acquired at defocus –3.5 and denoised using NAD filter with k value 5 and 5 iterations. Scale bar: 50 nm.

**Figure S1**

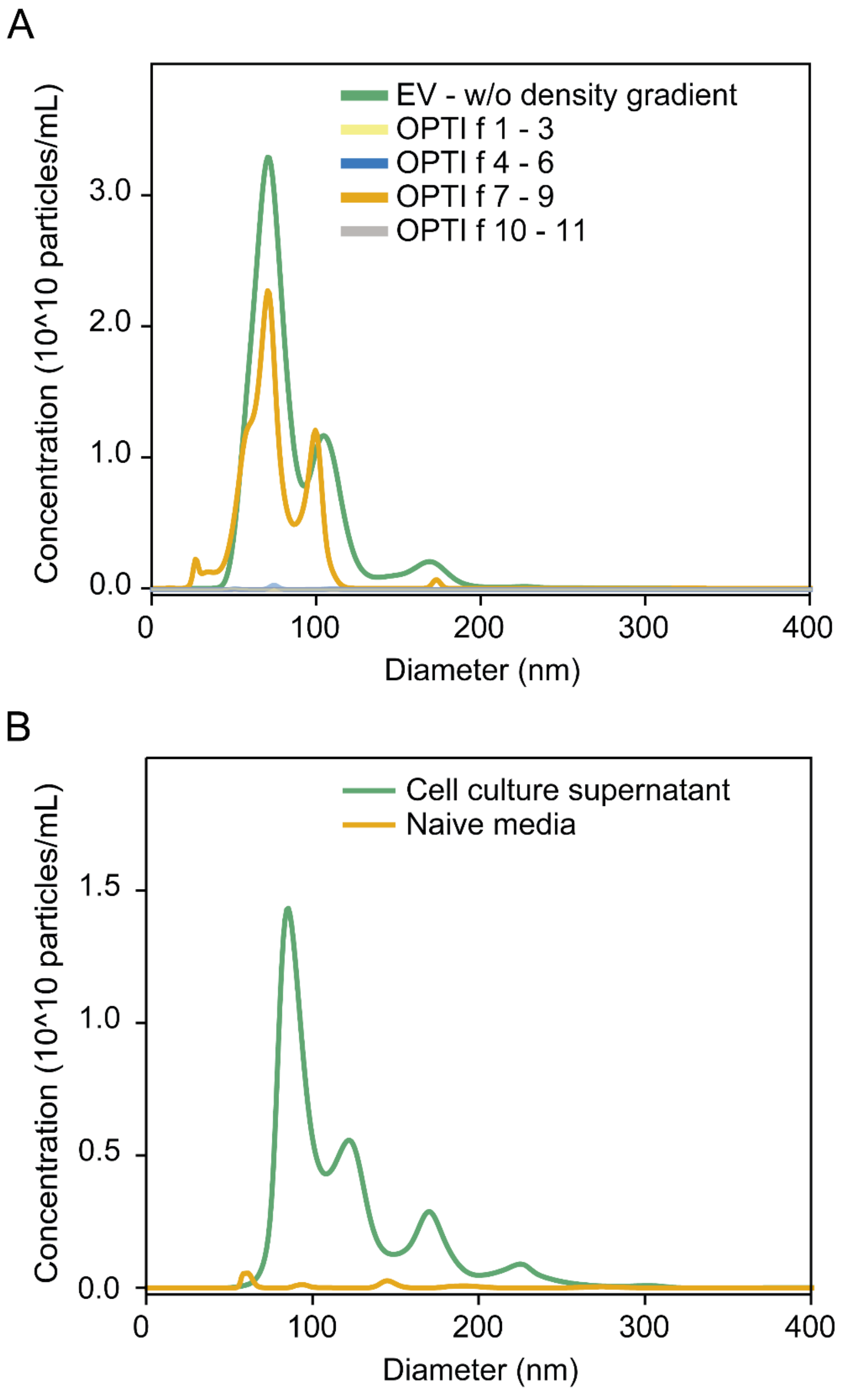
A
EV - w/o density gradient
OPTI f 1 - 3
OPTI f 4 - 6
OPTI f 7 - 9
OPTI f 10 - 11
Concentration (10^10 particles/mL)
0.0
1.0
2.0
3.0
0
100
200
300
400
Diameter (nm)
B
Cell culture supernatant
Naive media
Concentration (10^10 particles/mL)
0.0
0.5
1.0
1.5
0
100
200
300
400
Diameter (nm)

### Figure S2

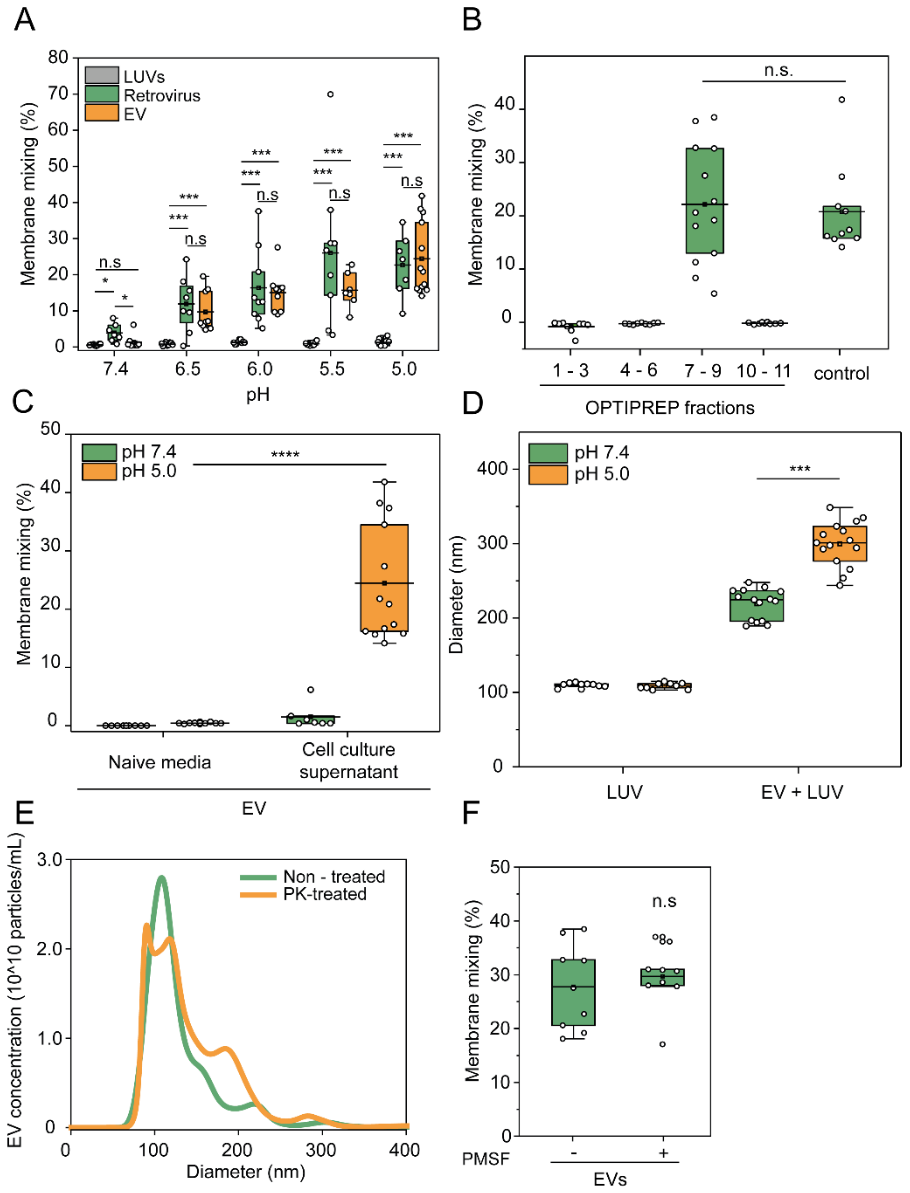
A
B
C
D
E
F
LUVs
Retrovirus
EV
Membrane mixing (%)
pH
7.4
6.5
6.0
5.5
5.0
n.s
***
*
OPTIPREP fractions
1 - 3
4 - 6
7 - 9
10 - 11
control
n.s.
pH 7.4
pH 5.0
****
Naive media
Cell culture supernatant
Diameter (nm)
LUV
EV + LUV
EV concentration (10^10 particles/mL)
Non - treated
PK-treated
PMSF
EVs

**Figure S3**

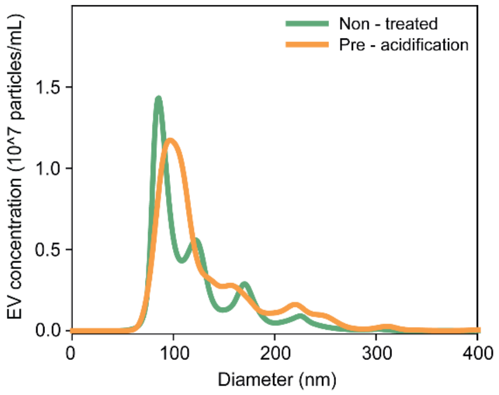
Non - treated
Pre - acidification
EV concentration (10^7 particles/mL)
0.0
0.5
1.0
1.5
0
100
200
300
400
Diameter (nm)

**Figure S4**

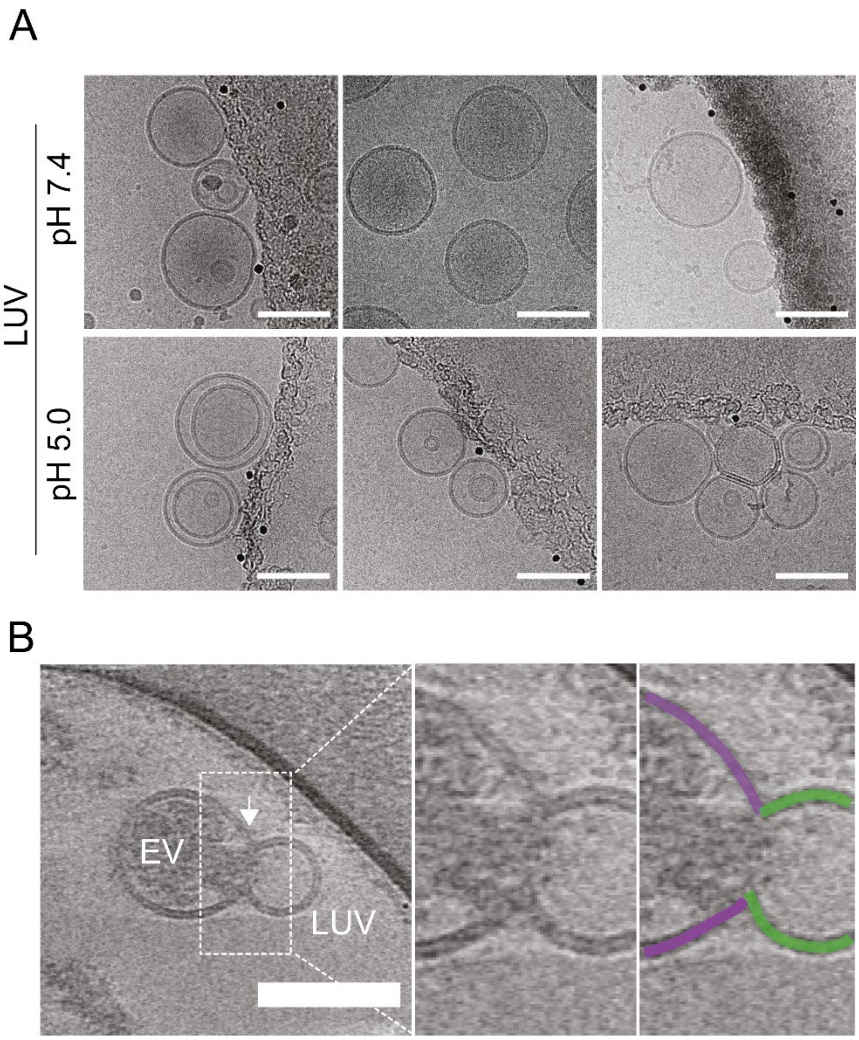
