## Supplementary figures and images for "Extracellular vesicle fusion visualized by cryo-EM"

### Original Gels of Western Blot figures

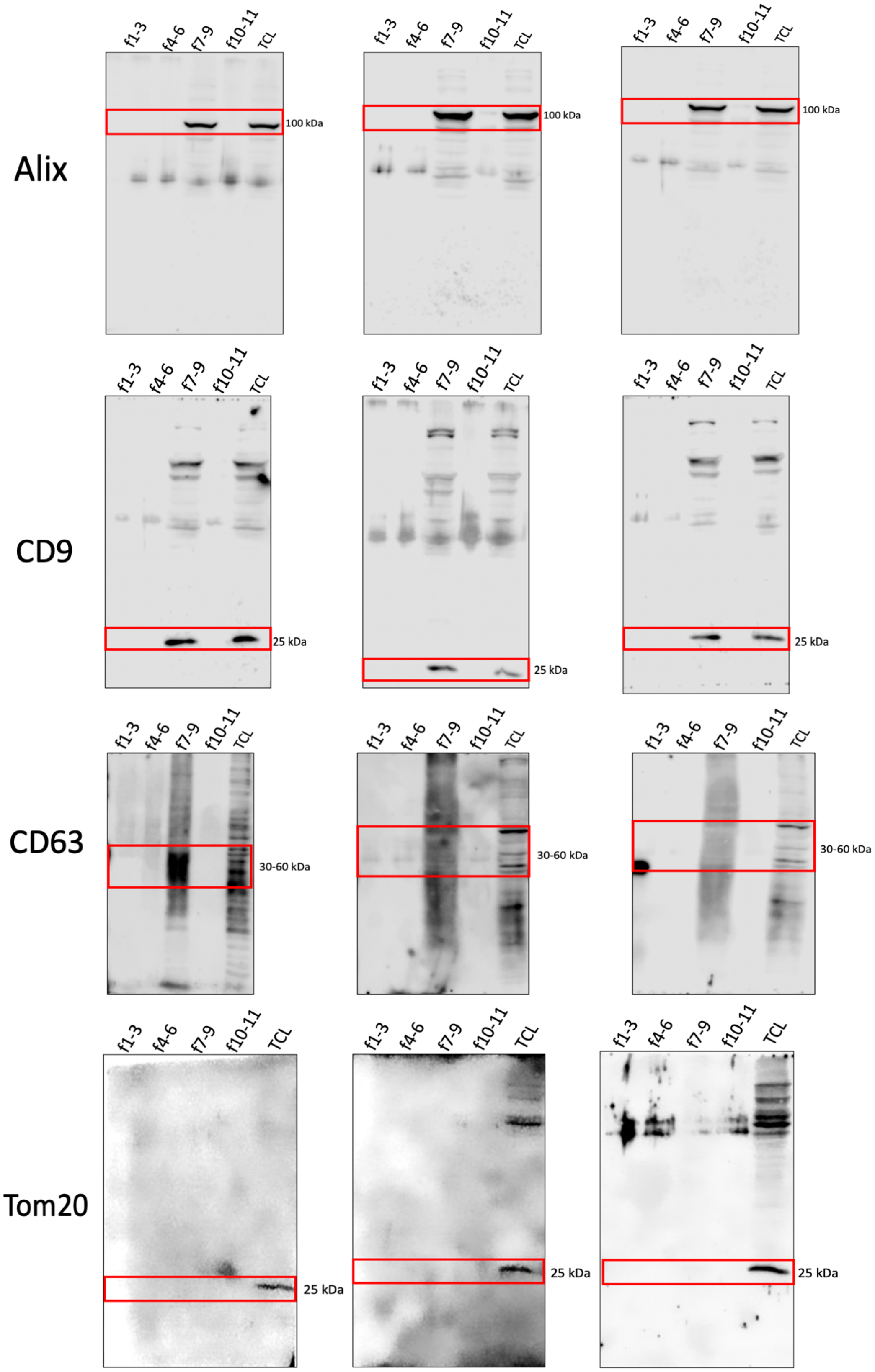

#### Fig. 2

Original membrane for Figure 2 F

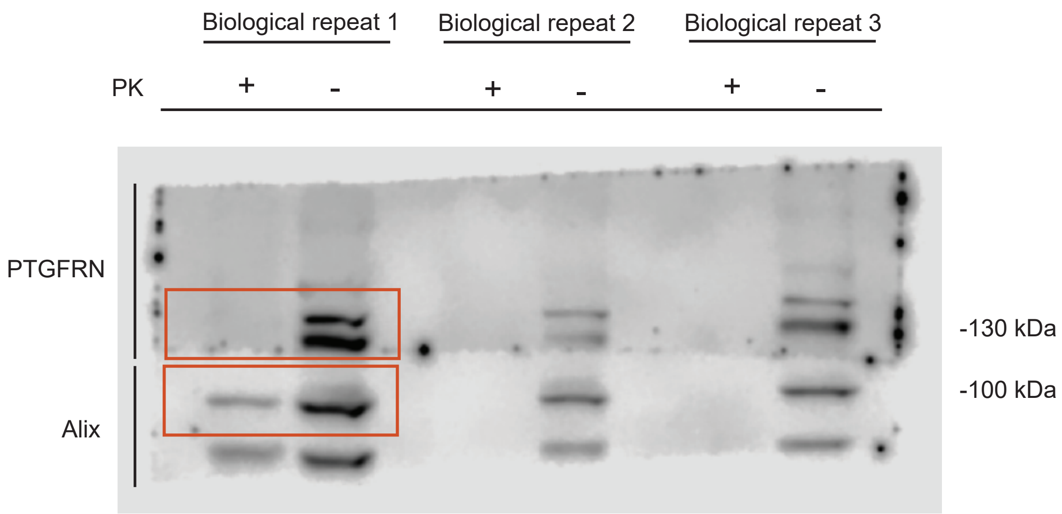
